# DAPTEV: Deep aptamer evolutionary modelling for COVID-19 drug design

**DOI:** 10.1101/2022.11.30.518473

**Authors:** Cameron Andress, Kalli Kappel, Miroslava Cuperlovic-Culf, Hongbin Yan, Yifeng Li

## Abstract

Typical drug discovery and development processes are costly, time consuming and often biased by expert opinion. Aptamers are short, single-stranded oligonucleotides (RNA/DNA) that bind to target proteins and other types of biomolecules. Compared with small-molecule drugs, aptamers can bind to their targets with high affinity (binding strength) and specificity (uniquely interacting with the target only). The conventional development process for aptamers utilizes a manual process known as Systematic Evolution of Ligands by Exponential Enrichment (SELEX), which is costly, slow, dependent on library choice and often produces aptamers that are not optimized. To address these challenges, in this research, we create an intelligent approach, named DAPTEV, for generating and evolving aptamer sequences to support aptamer-based drug discovery and development. Using the COVID-19 spike protein as a target, our computational results suggest that DAPTEV is able to produce structurally complex aptamers with strong binding affinities.

**Author summary:** Compared with small-molecule drugs, aptamer drugs are short RNAs/DNAs that can specifically bind to targets with high strength. With the interest of discovering novel aptamer drugs as an alternative to address the long-lasting COVID-19 pandemic, in this research, we developed an artificial intelligence (AI) framework for the in silico design of novel aptamer drugs that can prevent the SARS-CoV-2 virus from entering human cells. Our research is valuable as we explore a novel approach for the treatment of SARS-CoV-2 infection and the AI framework could be applied to address future health crises.

## Introduction

Viruses contain *deoxyribonucleic acid* (DNA) or *ribonucleic acid* (RNA) but are incapable of self-reproduction and rely on commandeering the cell’s protein creation capabilities to reproduce. After the viral protein has been successfully reproduced, it goes on to infect other cells in a process known as *viral proliferation*. Attaching and injecting of viral DNA or RNA to host cells is achieved through binding to cellular receptor through *receptor-binding domain* (RBD), an area on the viral protein evolved to specifically bind hosts’ cell receptor. In the case of the SARS-CoV-2 spike protein, the RBD targets the lung cell angiotensin-converting enzyme (ACE2) receptor [1–6].

Typical drug discovery focuses on either preventative triggering of host’s immune system or interrupting the life cycle of the virus. The former usually refers to vaccines. The latter is known as a therapeutic, attempting to halt the infection process in a currently-infected host [1, 7, 8]. While vaccines tend to focus on infection prevention [8, 9], therapy development can help alleviate suffering following infection.

*Aptamers* are short, single-stranded, oligonucleotides that can bind to targets with high affinity and specificity and it is hypothesize that specific aptamers can halt a virus during its life cycle by binding to the viral protein RBD there-by inhibiting its binding to host cell receptors. Aptamers can be created and modified easily and can bind to specific targets. One of the main ways to design aptamers is through a process known as *systematic evolution of ligands by exponential enrichment* (SELEX) [10] or *high-throughput SELEX* (HT-SELEX) [11] which applies high-throughput sequencing in each SELEX cycle. Design of aptamers through either SELEX or HT-SELEX is a slow, highly experimentally demanding process, possibly still providing sub-optimal designs [12–14] as these approaches rely of essentially a random search of top hits that are highly dependent on the initial choice of libraries [1, 8, 14–20].

*Artificial intelligence* (AI) techniques involve the modelling of brain intelligence to solve various challenging tasks [21]. As the most influential area in AI, *machine learning* (ML) improves intelligent agents or models using data and experience [22]. In the past few decades, machine learning has seen unprecedented progress. A variety of supervised and generative *deep learning* (DL) methods, *i.e*. neural networks, have been designed and achieved state-of-the-art performances in many domains [23]. Even though various AI and ML models have been recently developed for small-molecule drug design and achieved promising results [24–27], the end-to-end design of aptamers is a relatively unexplored domain [16, 19]. This is partially due to the unfamiliarity of aptamers in the AI community, unavailability of quality data, and the shortage of chemoinformatics tools for this class of molecules. In this research, to determine if AI approaches can accelerate aptamer drug discovery and development for the treatment of SARS-CoV-2 infection, we developed an *in silico* RNA aptamer design process, named *deep aptamer evolutionary model* (DAPTEV), similar to the experimental SELEX process. DAPTEV takes advantage of the embedding and generating capacity of *deep generative models* (DGMs), the high-throughput exploration power of evolutionary computation, and the quantitative measure of RNA-target binding affinities through molecular mechanism. The performance of DAPTEV was evaluated using the SARS-CoV-2 spike protein as a target. This work has the following major contributions: (1) we designed a novel end-to-end aptamer generation and optimization model to explore its performance, which is highly informative to researchers in the drug design domain; (2) using the SARS-CoV-2 spike protein as a target, we assembled a benchmark dataset for aptamer drug development, which can be used and improved by researchers in the future; (3) we applied our framework to search for RNA aptamer therapeutics for COVID-19 treatment, discovering new aptamers that can be further validated as an alternative means to address our pandemic challenge.

## Related Work

Until recently, patents for the SELEX process limited the potential exploration and innovation in the field of aptamer-based drug development [19]. Moreover, sources on ML applied to aptamers were even more scarce. However, with the patents now expired, new research is being released, some of which implement a model related to the one proposed in this research.

### Unsupervised Aptamer Identification Methods

A naive method for aptamer selection in SELEX is based on read counts. However, due to bias in *polymerase chain reaction* (PCR) [28], the high abundance of reads does not indicate a high binding affinity. To address this issue, early computational methods focused on aptamer identification from SELEX pools using *clustering* and *motif finding*. Clustering techniques, such as AptaCluster [29] and FASTAptamer-Cluster [30], attempt to learn aptamer sequence commonalities, but suffer from a lack of secondary structure considerations. Motif-finding approaches, such as MEMRIS [31], Aptamotif [32], MPBind [28], and APtaTrace [33], attempt to identify structure patterns with strong binding affinities. However, motif-finding approaches do not consider entire secondary structures. Both options struggle with processing times associated to large HT-SELEX libraries. To address these issues while utilizing the strengths of these two approaches, *Sequential Multidimensional Analysis AlgoRiThm for aptamer discovery* (SMART-Aptamer) is presented in [14] which accurately and efficiently identifies aptamers from SELEX libraries containing hundreds of millions of short sequences. SMART-Aptamer first applies a Markov clustering method to obtain aptamer families and then filters out the majority of the aptamers using multiple scores considering the enrichment of motifs, the abundance of aptamer families, and the overall secondary structures.

### Supervised Aptamer-Protein Interaction Prediction Methods

Aptamers that bind to a small number of protein targets have been selected to form limited aptamer-protein interaction (API) data, enabling the use of supervised models to learn from known APIs and then predict whether a new aptamer interacts with one of the protein targets listed in the training data. Due to limited training samples, conventional classifiers and hand-crafted features have been applied to API prediction and obtained moderate performance (70-80% accuracy). In [34] a *random forest* classifier learns on an API dataset containing a few hundred positive samples. Their input features include nucleotide composition, traditional amino acid composition, and pseudo amino acid composition. Similarly, in [35], an ensemble method (random forests) with hybrid features is presented to predict APIs using *Pseudo K-tuple Nucleotide Composition* (PseKNC) features to encode aptamers along with protein features including *discrete cosine transformation* (DCT), disorder information, and bi-gram *position specific scoring matrix* (PSSM). The task of ncRNA-protein interaction prediction is similar to API prediction, but has more data available to enable the training of a deep classifier. For instance in [36], RPITER, a hierarchical deep learning model with *convolutional neural network* (CNN) and *autoencoder* (AE) modules, is developed to automatically learn features from a few thousands ncRNA-protein pairs and obtained promising performance for ncRNA-protein interaction prediction. When more API data becomes available in the future, similar deep models are anticipated for API prediction.

### Aptamer Generation

Im *et al*. used a generative model to build statically sized (20 nt) DNA and RNA sequences that bind to a target protein [37]. At the time of publication, it was specified that their research was ongoing, but they were able to train a *long short-term memory* (LSTM) model on a “huge dataset of sequences from high-throughput experimental technologies … such as HT-SELEX or CLIP-seq”. This dataset, which is still not available publicly, was obtained from DeepBind [38]. DeepBind was used to estimate the affinity and specificity of generated sequences. The target proteins in their research were as follows: DRGX, GCM1, OLIG1, RXRB, NFATC1, NFKB1, and MBNL1. It was found that the produced sequences possessed structural motifs similar to known motifs, and that the produced sequences had a strong binding affinity and specificity to their intended target [37]. This research was continued by Park *et al*. and a similar conclusion was drawn for RNA aptamers specifically [39]. In [40], Iwano *et al*. propose RaptGen which is similar to [37] and [39]. They too utilized DeepBind, however, RaptGen implemented a CNN-LSTM as their encoder and a profile *hidden Markov model* as their decoder. Additionally, while RaptGen starts with static sequence lengths (10 nt), it utilizes some post-processing techniques to extend the sequence lengths to a set of other fixed sizes, technically achieving sequence generation with variable lengths. Iwano *et al.’s* target proteins were the TGM2 and *αvβ*3. The main difference between the research mentioned above and our own is as follows: (1) our model allows for the specification of a target RBD and calculating binding affinity based on thermodynamic principles rather than estimating binding via ML surrogate models; (2) our model optimizes aptamers to produce sequences with better binding affinity and specificity to the intended target instead of only generating aptamers having the same distribution as the training data; and (3) our model allows for true variable length sequences as input and output rather than fixed sized sequences.

In [13], Lee *et al*. present an aptamer generation method called Apt-MCTS. This method uses limited positive and negative aptamer-protein pairs to train a random forest classifier as a surrogate model to predict whether an aptamer can bind to a fixed protein target (one of 6GOF, 3V79_1, 5VOE, 3SN6_4, 2RH1_1, and 1ERK_1). The prediction result is used as a score to guide a *Monte Carlo tree search* (MCTS) [41] process for the generation of new aptamers. Apt-MCTS and our DAPTEV research utilize the same starting real aptamer data, but we further expanded this data by generating additional RNA sequences for model training. Apt-MCTS requires the availability of aptamer-protein pairs to train a surrogate model for API predictions. This disables the application of Apt-MCTS in our situation where experimental aptamer-spike protein interaction data are unavailable.

Wornow proposed a conditional VAE to generate novel strong-binding aptamers [19]. This too is similar to our work and findings. However, Wornow’s work utilized 8 rounds of SELEX data as a proof of concept on DNA data only. This creates a limitation of requiring experimental data to be collected before the model’s efficacy can be illustrated or be used in practice and does not explore the RNA landscape. The target for Wornow’s research is the chemotherapeutic agent daunomycin. Binding affinity was approximated based on ground truth fitness scores which were later confirmed by wet-lab experiments. They avoid the use of *molecular dynamics* (MD) simulations, stating they are computationally expensive and infeasible for use in the field.

### Molecular Optimization

Grantham *et al*. have developed a multi-objective deep learning framework named *deep evolutionary learning* (DEL) for small-molecule design. DEL is beneficial for improving sample populations in terms of property distributions and outperforms other multi-objective baseline molecular optimization algorithms [26]. While DEL does not work with aptamer data, it does focus on molecular optimization. Thus, it serves as inspiration for our DAPTEV.

## Method

Our *deep aptamer evolutionary modelling* (DAPTEV) framework is visualized in Fig. 1. It is a hybrid approach that integrates the strengths of DGMs (variational autoencoders) for aptamer encoding and modelling, computational intelligence (evolutionary computation) for aptamer optimization, and bioinformatics tools for RNA secondary and tertiary structure prediction and RNA-protein folding-and-docking (Rosetta). The continuous docking score is used as the fitness value or objective in the evolutionary computation component to guide aptamer optimization.

**Figure 1.**
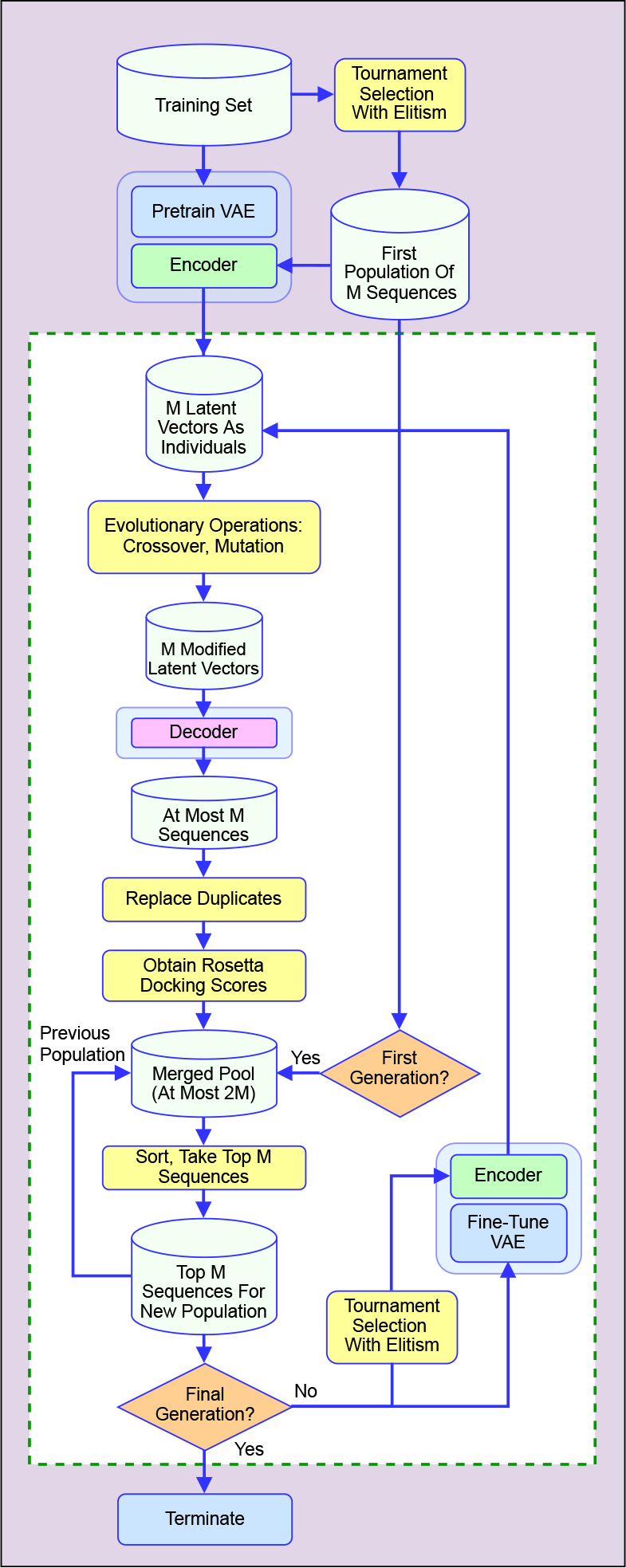
Flowchart depicting the DAPTEV process for aptamer design.

First, a dataset must be collected for the training of the deep learning model and the initialization of the optimization process. The dataset should include a large number of aptamer sequences of certain lengths, their corresponding secondary structures, and docking scores to a protein target of interest. The secondary structures are needed by the Rosetta package [42] for docking. If secondary structures are unknown, they can be quickly computed using Arnie [43]. For details of assembling data to generate aptamers targeting SARS-CoV-2 spike protein, please refer to Section.

As an initialization step, a *variational autoencoder* (VAE) [44] is pretrained using the dataset. This VAE provides an encoding process that transforms an aptamer sequence to a vector of continuous values which is called the *embedding vector* or *latent vector* of the aptamer. Since the latent space is continuous, various metaheuristic techniques can be applied in this space for aptamer optimization. The decoder of the VAE enables a decoding process that converts a latent vector (either randomly sampled or modified using computational intelligence operations) to an aptamer sequence. Please refer to Section for details of our developed VAE.

The main body of the DAPTEV framework is an iterative evolutionary process. At the beginning of the loop, a population of aptamers are randomly sampled from the original dataset, then they are passed to the encoder to obtain latent representations. Evolutionary operations are applied to these latent representations and then these modified latent representations are decoded using the decoder to obtain their corresponding primary sequences. The fitness of these new sequences can be measured through aptamer-protein docking. After that, the newly produced aptamer sequences and the previous population of aptamer sequences are ranked together and only top aptamers are kept to form a new population of aptamers for the next generation. In the next generation, this new population is used to fine-tune the VAE model and then uses the encoder of the fine-tuned VAE to project the new population into the latent space where evolutionary operations are again applied. See Section for detailed discussion of the evolutionary operations. Each time when a new aptamer sequence is generated, Rosetta needs to be called to calculate the docking score. The docking component is detailed in Section. Furthermore, docking is a time-consuming process. Its speed-up is discussed in Section.

### Variational Autoencoder for Aptamer Modelling

VAE is a probabilistic deep generative model that takes advantage of the autoencoder architecture where the encoder network corresponds to the inference *q_ϕ_*(***z***|***x***) and the decoder network realizes the generative component *p_θ_*(***z***)*pθ*(***x***|***z***). Here ***x*** is the vector of random variables for observed data (i.e., an aptamer sequence in our case), ***z*** is the vector of latent random variables, ***ϕ*** and ***θ*** are respectively the parameters of the inference network (encoder) and generative network (decoder) [44]. In our work, since aptamers are represented in primary sequences, *gated recurrent units* (GRU) [45] are used in the encoding and decoding processes. Thus, the resulted VAE is a probabilistic Seq2Sep structure. See Fig. 2 for a depiction of the implemented VAE.

**Figure 2.**
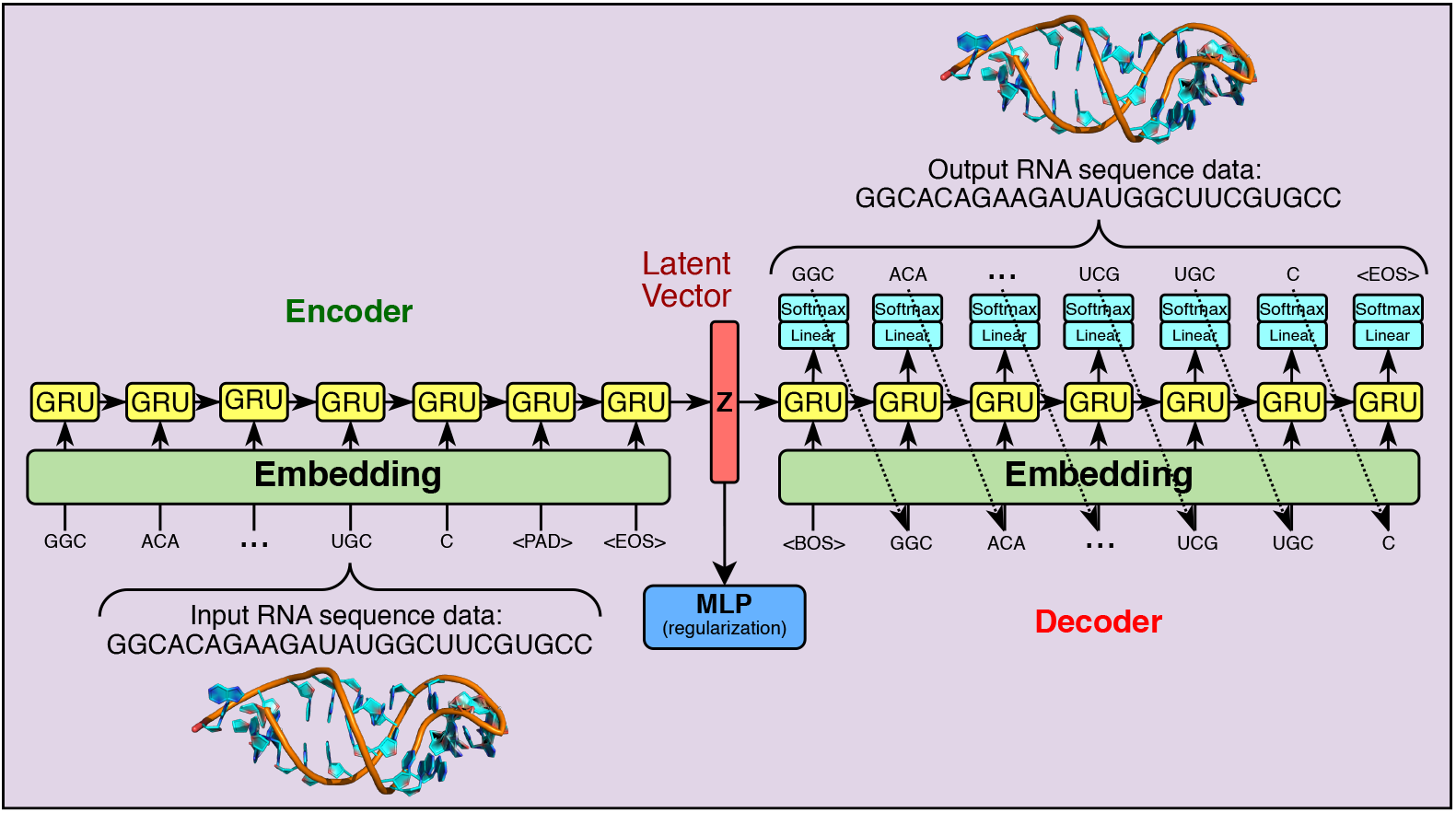
Depiction of the implemented VAE.

Using “A” for adenine, “C” for cytosine, “G” for guanine, and “U” for uracil, an aptamer sequence of length *L* can be represented as ***x*** = [*x_i_*|*x_i_* ∈ {*A, C, G, U*}, *i* = 1, 2,…, *L*]. The *word embedding* representation techniques [46, 47] can be used here by treating each letter as a word. However, this alone does not provide the model with enough context for how an RNA structure will interact with itself and the possible structural motifs that can occur. Thus, the combination of all possible 1, 2, and 3-mer strings (as tokens) are included in the vocabulary 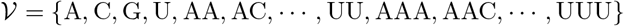. Conversely, adding too many combinations could drastically increase the computing time and the need for more training data. Thus, we restrict our vocabulary up to 3-mer. Three more special tokens are inserted into the vocabulary: <BOS> for beginning of sequence, <EOS> for end of sequence, and <PAD> for padding. Both the <BOS> and the <EOS> tokens inform DAPTEV of the character boundaries for each sequence. When working with sequences, one inevitably must choose how to handle sequences of variable lengths. Rather than restricting all sequences to the same length, limiting the versatility and learning capabilities of the model, <PAD> tokens are inserted into each sequence. In the encoding process, the sequence needs to be parsed into tokens. The 3-mer tokens have the highest priority in this process. A 2 or 1-mer token only appears at the end of the parsed result when a 3-mer parsing is no longer possible. Taking sequence GGCACAGAAGAUAUGGCUUCGUGCC for example, the parsed sequence of tokens is [GGC, ACA, GAA, GAU, AUG, GCU, UCG, UGC, C]. The decoding process follows a similar scheme. The decoding will stop when (1) it generates a 2 or 1-mer token, (2) an <EOS> token is generated, or (3) the length of the generated sequence meets the prespecified maximal length.

The bottleneck between the encoder and decoder is the latent layer representing the embedding of the whole sequence. From the autoencoder perspective, a latent vector contains the compressed key information about the corresponding input and this latent vector can be decoded to reconstruct the input. From the generative model perspective, the latent vector, following a simple prior distribution (a multivariate standard Gaussian distribution in our case), is the start of the generation process. The feasible domain of the latent vector forms the latent space.

Up to this point, the VAE designed above is not necessarily learning what makes an RNA sequence good or bad with respect to the Rosetta docking score. The VAE should be able to produce sequences with strong docking capabilities to a specified target. It is for this reason that an MLP component was included in the architecture of the VAE. This MLP performs score-based classification and further regularizes the latent representation. It takes a latent vector as input and predicts whether the sequence associated with the latent vector is good in terms of target binding, this vector and its associated score will be given to the MLP. A lower docking score implies a tighter binding. Thus, a “good docking score” is defined as ≤ the user-chosen score threshold parameter and a “bad docking score” is defined as > the score threshold. This converts the scores to 1 (good) or 0 (bad). Note that one cannot simply set the score threshold to any low value desired as it can result in highly imbalanced data. To determine the optimal score threshold, one may locate the lowest score that still labels at least 25% of the data to Class 1 (as a rule of thumb only). For example, it was found that 26.28% of our data (see Section) fell within the docking score range of 3,500 and below for this research. Thus, a score threshold of 3,500 was chosen in our experiment. Why is the docking quality prediction modelled as a classification problem rather than a regression problem? According to our experience in docking score modelling, regression usually has unsatisfactory performance.

The loss function of the VAE to be minimized in DAPTEV is formulated below

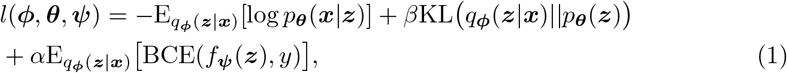

where the first term corresponds to the reconstruction error, the second term is the *Kullback-Leibler* (KL) divergence that pushes the posterior towards the prior, and the last term is the *binary cross-entropy* (BCE) loss for the MLP (parameterized by ***ψ***) performance where *f_ψ_*(***z***) is the predicted value and *y* is the actual class label. The trade-off hyperparameters *α* and *β* control the balance among these three terms. The KL loss term tends to vanish easily during training in the vanilla VAE where *β* = 1. This can negatively affect the reconstruction learning process. To address this issue, we implemented a technique known as *KL annealing* [26, 48] to control the value of *β*. The annealing process applies a small weight, between 0 and 1, to the KL loss value and linearly increases this weight during training.

### Evolutionary Operations for Aptamer Optimization

Each evolutionary generation ensures the VAE has finished a round of training (fine-tuning) such that DAPTEV can then begin the sequence optimization step. This is when DAPTEV searches for better sequences and update its running data accordingly in every generation. Three Darwinian evolutionary operations are involved in this step.

The first operation is known as *tournament selection with elitism. Elitism* is the action of selecting the top e best individuals (sequences) in a population to be carried forward into the next generation. This ensures the best individuals remain in the running data and are not filtered out during tournament selection. Tournament selection is the process of randomly sampling sequences from the population and having these candidates compete in a tournament. The winner of each tournament will be selected as a *parent* to produce the next generation of *children*. This process is repeated multiple times until the parent pool equals the chosen population size. The tournament selection operation is controlled by two hyperparameters. One is the *k* value which indicates how many contenders with which to build a tournament round. The higher the k value, the more likely a stronger performing individual will be selected to win the tournament and proceed as a parent for the next generation. This, in turn, means there is a higher likelihood that poor-performing individuals will not be selected to proceed as parents (referred to as *selection pressure*), which can often have a detrimental effect on exploration. The other hyperparameter is the *selection rate*. This parameter creates a possibility for the best-performing individual in a tournament pool to opt out, allowing some poor-performing individuals to be selected instead.

The second evolutionary operation is *crossover*. This is the process of producing children from two parents, attempting to simulate a child obtaining features from their parents. After the selection operation is performed, the sequences are passed through the VAE’s encoder to obtain their latent representations (***z**s*). Then, the crossover operation is applied to these parent ***z*** vectors whereby parent ***z**_pa_* and parent ***z**_pb_* will each copy random parts of their latent vectors to new children vectors ***z**_ca_* and ***z**_cb_*. This procedure is repeated until the number of children equals the population size. Similar to the tournament selection, this operation also has a probability of occurring (called *crossover rate*). Note that too small of a population size will result in a low amount of *“genetic diversity*” which can quickly converge to a local minimum (suboptimal solution). This means that there will not be enough sequences in a given generation to produce sufficiently dissimilar children from the parent sequences. Eventually, all produced sequences could start to resemble each other and/or the algorithm will be unable to find better sequences due to the lack of “genetic diversity” among available sequences.

The last evolutionary operation is *mutation*, mimicking random mutations in evolution. Some random influence should be allowed such that an individual may find themselves performing significantly better than the rest. In DAPTEV, mutation is implemented by selecting, with a probability called *mutation rate*, a child from the crossover result. Then a random index of that child’s ***z*** vector is chosen and replaced with a random, normally distributed value. The mutation rate should be very small. A large rate will result in the algorithm essentially performing just a random search, corrupting any learning.

Once the new latent vectors have completed a round of evolutionary operations, they are sent through the VAE’s decoder. This will generate the sequences based on the VAE’s previously trained decoding capabilities. The output will be a list of new sequences. Any duplicates in this list, as compared to themselves, the starting data, and any previously predicted sequences will be removed and replaced with new, folded, random sequences. However, docking scores for these new sequences are unknown. Thus, the docking simulation is repeated before the VAE continues to fine-tune using the new population.

### RNA-Protein Docking

Rosetta models RNA-protein complexes by simultaneously folding and docking an RNA to a protein surface via a statistical RNA-protein docking scoring function [42]. This allows one to specify a target protein and the target’s RBD. There are two scoring functions within DAPTEV’s usage of Rosetta: *native Rosetta docking score function* and *constraint scoring function*.

The native Rosetta scoring function is a low-resolution, coarse-grained, knowledge-based (statistical) RNA-protein potential. This serves as an energy function for scoring Monte Carlo steps within the tertiary structure prediction and docking simulation. The lower the returned score is after the tertiary structure prediction and docking simulation, the more stable the predicted complex is considered to be. In Rosetta, all previously published score terms describing RNA structure and RNA-protein interactions are included in this scoring function [42], while also providing rapid computation and maintaining coarse-granularity. It is reported in [42] that, over ten popular scoring systems, the best-performing scoring models achieved an average atomic *root-mean-squared deviation* (RMSD) of 11.6 A for 3dRPC and 10.2 Å for DARS-RNP, whereas Rosetta’s score function achieved an RMSD of 6.4 ^Å^.

The constraint scoring function applies to where on the protein the RNA docks. This constraint function allows one to specify the target’s RBD region without having to force a fixed binary interaction between specific atoms of the RNA and the target protein. Instead, one indicates to Rosetta their chosen constraint type, to which atoms on the target and the RNA the constraint applies, and what built-in formula the constraint will use to calculate an energetic penalty. This penalty will be applied against the returned Rosetta score to passively discourage the RNA from docking elsewhere on the target. As a result, the RNA has a range of acceptable distances it can deviate from the target’s RBD during the docking simulation.

DAPTEV uses an “atom pair” constraint with a flat harmonic function. The middle position (nucleotide) of every produced RNA sequence is chosen as the constrained RNA nucleotide. This function is formulated below,

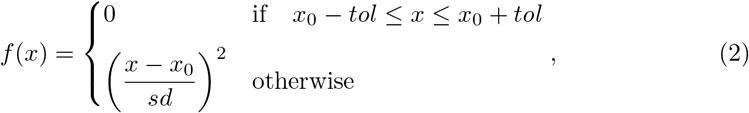

where, *x* represents the distance between the two atoms (this varies as Rosetta attempts multiple conformations), *x*_0_ is the ideal distance between the atoms, *sd* is the standard deviation from the ideal distance, and *f*(*x*) is the returned penalty. This function produces a penalty of 0 if *x* falls in the range [*x*_0_ – *tol*, *x*_0_ + *tol*], where *tol* means tolerance. In our research, we set *x*_0_ = 0, *sd* = 0.125, and *tol* = 1.

### Computing Time Improvement

A challenge for utilizing Rosetta in DAPTEV is the computational demand and time required to perform docking simulations. For an experiment with population size 800, generation number 10, and 3 runs, it could take about 4 years if the *full* SARS-CoV-2 spike protein was used and docking simulations ran *sequentially*. The running time can be reduced to roughly one-third of a year by reducing the protein file to just the RBD. However, this is still too long to wait for one experiment to finish. As such, *multiprocessing* was implemented into DAPTEV. Multiprocessing is a technique to run multiple docking simulations on a computer’s multiple CPUs at the same time (in parallel). DAPTEV utilized 18 CPUs to effectively reduce computing time for each docking simulation from roughly 6.5 minutes (also using the reduced protein) to 1 minute per aptamer sequence.

## Experimental Results

### Data

In our experiments, we used the SARS-CoV-2 spike glycoprotein (closed state, PDB: 6VXX) [5] as the docking target with an RBD between residues 333 and 524 [2, 4–6, 49]. See Fig. 3 for an illustration. As it shows, the SARS-CoV-2 spike protein’s RBD represents a relatively small area at the apical face of the protein. Using the entire spike protein in the docking procedure would drastically increase the computation time. For example, performing five runs of the docking process with an RNA base count of 25 on the full SARS-CoV-2 spike protein took over 98 minutes to finish computing. Thus, one should remove all unnecessary residues from the target PDB file and save the new PDB structure and the new FASTA sequence. Unnecessary residues are those on the target that an RNA could not possibly interact with during the RBD-docking process due to constraint specification. Once these residues are removed, the PDB file must be cleaned (renumbered and sequenced).

**Figure 3.**
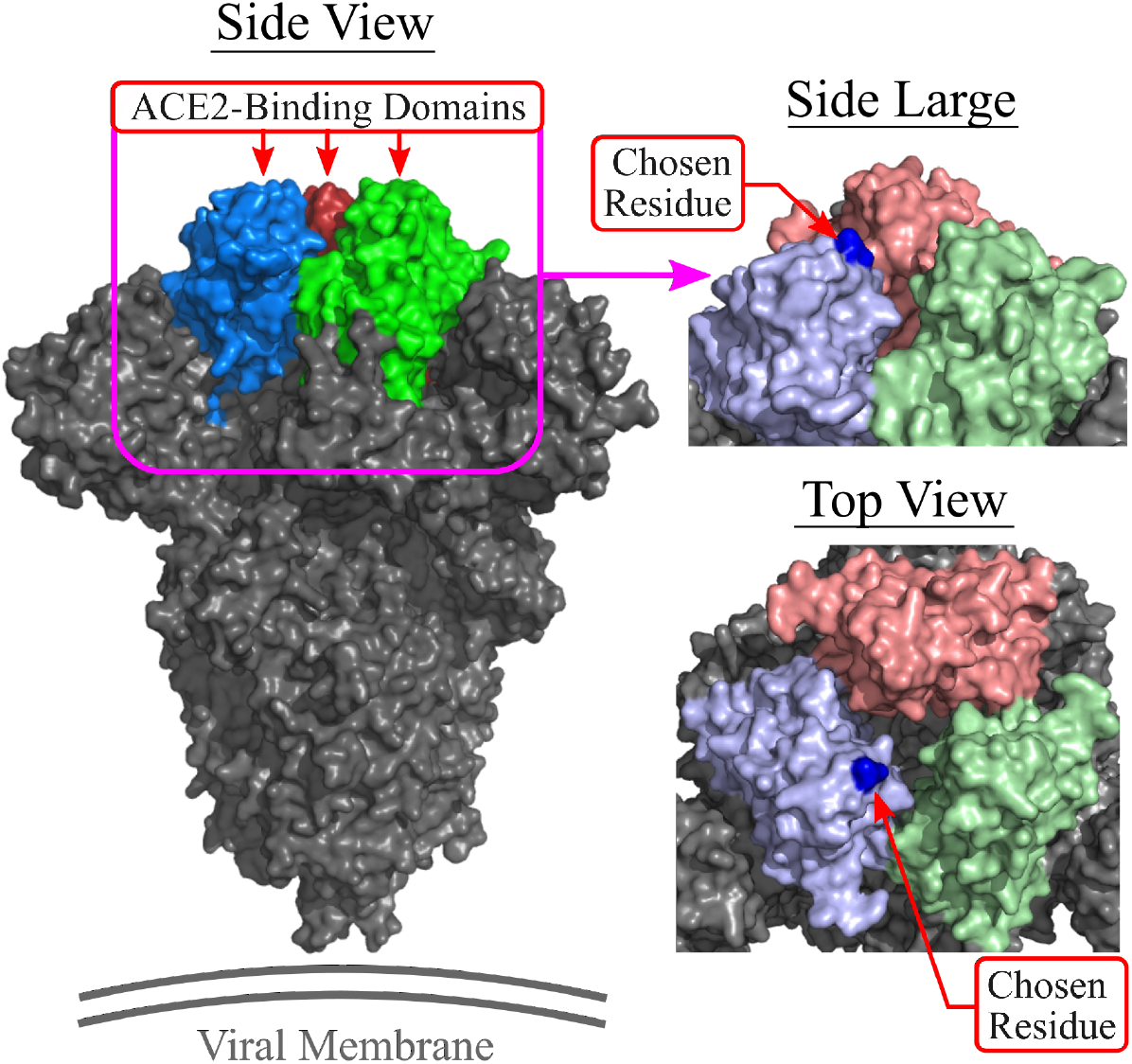
Location of the chosen residue (201A, atom C5, dark blue dot) on the renumbered SARS-CoV-2 spike protein RBD. Note: the full spike protein shown (left) is for reference but the protein file in DAPTEV is cropped. PDB: 6VXX.

For the implementation in the docking, the protein is cropped according to the RBD and renumbered. The chosen RBD residue number is 201 and the associated chain is “A” for the Rosetta constraint. Fig. 3 shows a visual of the chosen residue. The carbon atom “C5” is ubiquitous among RNA bases [50]. This was the reason behind choosing “C5” as the RNA atom parameter. Doing so required no additional calculations to be performed and was simple to implement.

While an entire dataset can be created using the random sequence generator, it is best to augment the dataset with known aptamers. Providing existing aptamer data will add diversity into the dataset, will expose the model to physical characteristics of existing aptamers, and will help the model learn these traits. This will also allow DAPTEV to rule out or confirm existing aptamers as potential solutions to the given problem as it is possible that a known aptamer is already a good fit for the specified target. As a result, supplying existing aptamer data could provide DAPTEV with a strong starting position before it begins exploring other options.

We do not allow unfolded RNA secondary structures to be created when performing random sequence generation. Associated secondary structures will be restricted to having at least one set of brackets (base pair connections). It has been observed during the experimental phase that unfolded secondary structures produce RNA tertiary structures that are more malleable when docking to the target’s RBD. Thus, unfolded sequences can better form to the target’s RBD topology, producing strong binding scores (binding affinity). However, aptamers are designed not only for high binding affinity but also for target specificity [1, 8, 15–18, 20]. An unfolded structure would not be considered “specific” to a target because it could potentially bind just as well to another target. Furthermore, the Rosetta scoring function for the docking process includes the quality of the RNA tertiary structure prediction. This means that the returned score for any sequence in this model may be deceptively “improved” due to encountering few stability-related penalties during the structure prediction and docking process. As a result, the scores produced by unfolded RNA structures are artificially and significantly better than their more structurally-complex peers. This can result in the algorithm producing many unfolded RNA aptamers, which would not be ideal for aptamer drug development as the goal is to optimize the aptamer discovery process for a specific target only (specificity).

After obtaining 849 unique known aptamer sequences [13, 51, 52], only 390 met the condition of being between the minimum (20) and maximum (40) sequence lengths. Of those 390 aptamer, 344 contained at least one connection (base pairing) in their secondary structures. 44 sequences were too small, 417 sequences were too large and an additional 44 sequences were unfolded (based on the returned Arnie secondary structure predictions). As 12,000 data points were required, an additional 11,656 random sequences were generated and scored. These sequences were restricted to having lengths between 20 and 40 nts, an approximate GC percentage of 50%, and containing at least one secondary structure connection.

### Hyperparameter Setting

In our experiments, the settings of hyperparameters for the data collection process, the evolutionary optimization process, and the structure and training of the VAE are listed in Table 1.

**Table 1.**
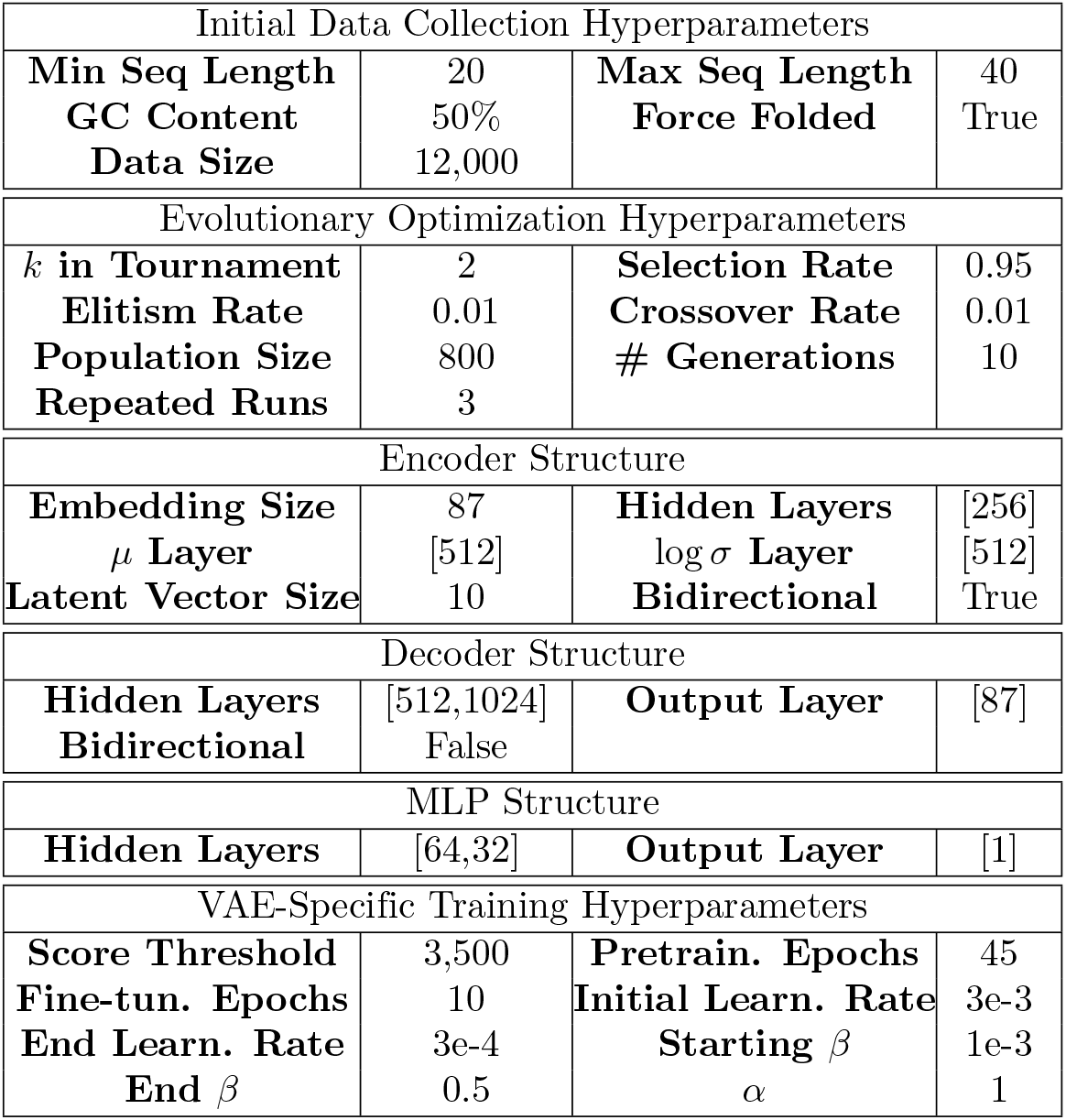
DAPTEV hyperparameters used in our experiments.

### Baselines

The experiment of DAPTEV is accompanied by two additional comparison models: a genetic algorithm on the sequence space (GA), and a “hill climer” algorithm (HC).

The GA baseline is to simply remove the VAE from DAPTEV, resulting in a GA operating on the problem space rather than the latent space, to see how their performance differs. It is worth noting that the VAE and the GA are not restricted to producing only folded structures. As such, these models do have a possibility of generating unfolded sequences. This is allowed to more deeply explore the solution landscape and potentially encounter stronger sequences through the prediction process. For the crossover function in GA, instead of selecting random indices of parent ***z**_pa_* and parent ***z**_pb_* (latent vectors), parents ***x**_pa_* and ***x**_pb_* will have random indices of the sequence space selected for crossover. The resulting children will be two new sequences. For the mutation function in GA, an individual is selected with a small probability. If selected, a random index of within the sequence is randomly chosen such that the corresponding letter is replaced with a random token from the vocabulary list excluding <BOS>, <PAD>, and <EOS>.

The HC baseline generates *random* sequences using our dataset creation script with the same chosen hyperparameters as the DAPTEV. This process is repeated for the same number of sequences, generations, and runs as the DAPTEV. The best performing sequences are carried forward into the next generation and all other sequences up to the chosen population size are discarded. This is known as a “hill climber” because this algorithm employs the simple heuristic of “always take the best” without any actual “learning”.

### VAE Performance

Fig. 4 shows how the VAE’s encoder, MLP, and decoder modules performed per epoch and over each generation (plotted separately on the same graph). The total loss values are also shown in this figure. In the first generation, the VAE is trained on the entire dataset of 12,000 sequences. It shows that, the KL loss can quickly decrease to a small value after 10 epochs. Each subsequent generation shows the encoder module continues to improve on the KL loss and approaches a near zero loss value. The MLP regularization module performs similarly. It shows that this module is also refining its performance. In fact, the module learned to perfectly classify the data halfway through generation five. The reconstruction loss graph suggests that, after the first generation, the decoder is learning how to reproduce the encoded latent vectors very well. The total loss graph shows the VAE’s overall performance per epoch per generation. As each model effectively reduced its loss values, it stands to reason that the total loss would reflect this performance.

**Figure 4.**
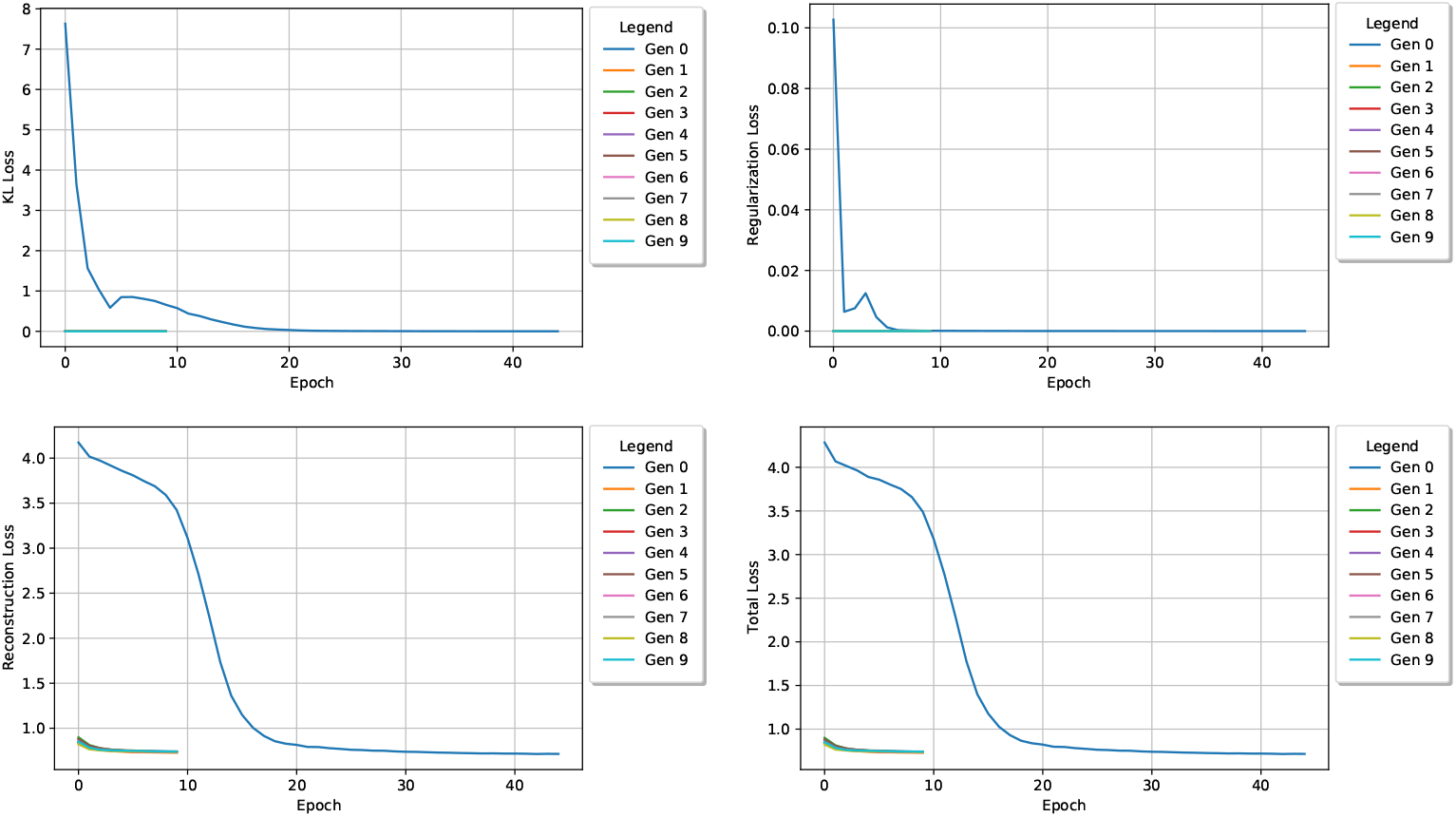
KL loss values (top-left), MLP regularization loss values (top-right), reconstruction loss values (bottom-left), and total loss values (bottom-right) per epoch for each generation. Note: total loss values include the KL weight factor over time.

### Comparison in Terms of Docking Score

Table 2 provides details on the overall docking score optimization performance for each model. Here, it can be seen that both DAPTEV and the GA improved significantly from their worst scores to their best scores, with GA outperforming DAPTEV. However, in comparison to the hill climber model, both DAPTEV and GA have performed better.

**Table 2.** The average score performance in the first and last generations for each experiment. Standard deviation provided in brackets. DAPTEV and GA were, respectively, run for 3 times. HC was simulated based on our original dataset, thus was only run once.

| Docking Score |  | DAPTEV | GA | HC |
| --- | --- | --- | --- | --- |
| First Gen | *Best* | 118 (14) | 100 (41) | 18,250 |
|  | *Mean* | 897 (207) | 781 (337) | 23,635 |
|  | *Median* | 925 (204) | 801 (343) | 21,957 |
|  | *Worst* | 1,256 (297) | 1,098 (489) | 60,677 |
| Last Gen | *Best* | 72 (31) | 32 (24) | 128 |
|  | *Mean* | 578 (49) | 292 (58) | 1,098 |
|  | *Median* | 602 (49) | 303 (59) | 1,122 |
|  | *Worst* | 796 (77) | 390 (86) | 1,530 |

Fig. 5 shows the best, worst, mean, and median performing docking scores produced by each model per generation. The provided graphs are scaled logarithmically as the hill climber (“Random” in the legend) experiment performed significantly worse than DAPTEV and GA. It is clear that GA produced better results than DAPTEV in terms of docking scores. The lowest (best) score produced by DAPTEV was 98 in generation three, whereas GA obtained its lowest score of 4 in generation nine. DAPTEV’s worst scores fell from 1,470 to 857 whereas GA’s worst score fell from 1,439 to 460. Both the means and medians seem to follow a fairly similar trend and are comparable in score output.

**Figure 5.**
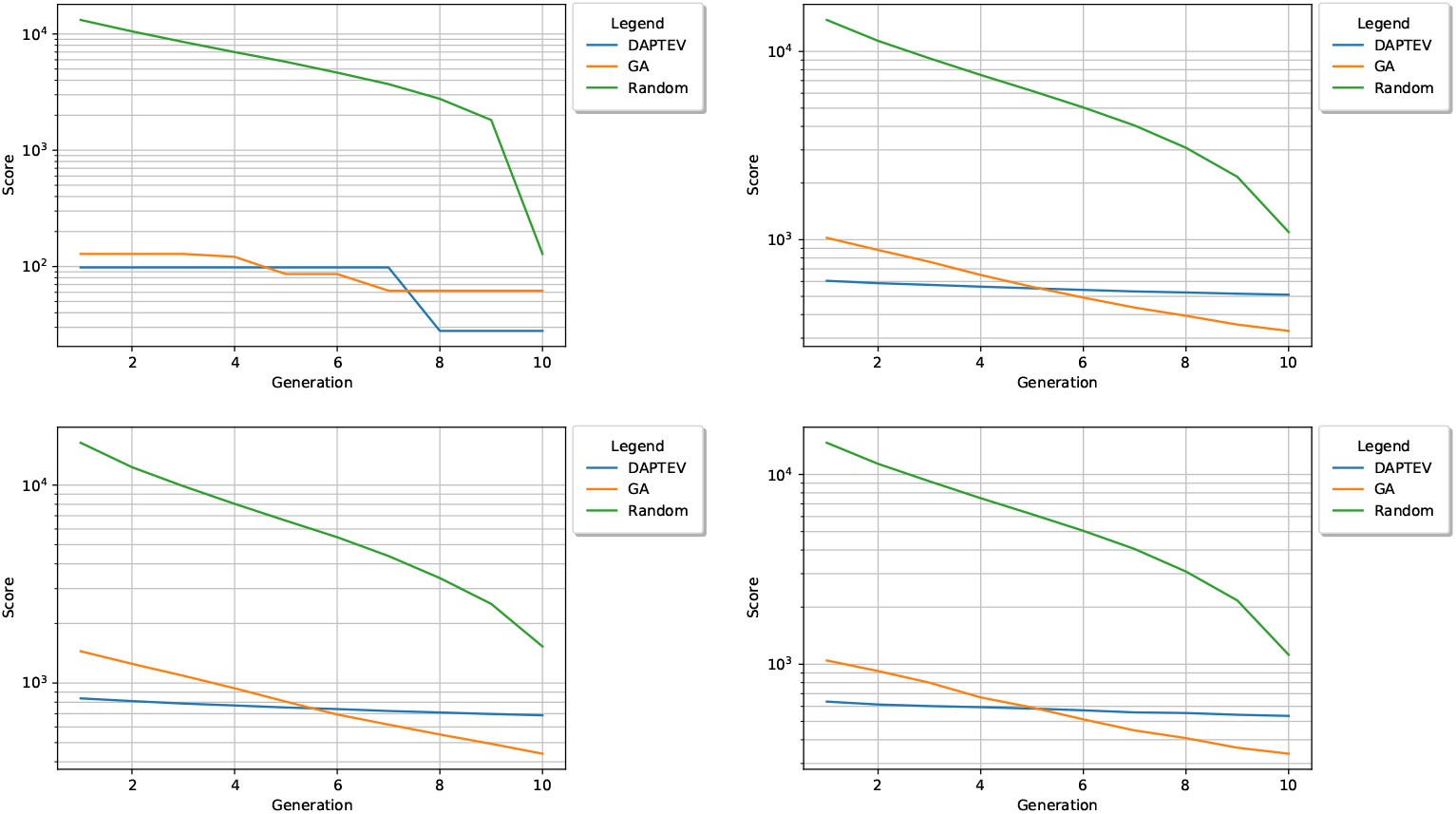
The three models’ best (top-left), worst (bottom-left), mean (top-right), and median (bottom-right) scores plotted per generation on a logarithmic scale in a computational run.

It also seemed prudent to consider the best five and worst five mean scores for each experiment. The absolute best and worst can send an extreme message and may not convey the true performance of each experiment. This is because the absolute best and worst do not include neighbouring sequence performance. If these models are indeed performing well, then one would expect to see the sequences near the best and worst improving overall. Fig. 6 depicts the best and worst five score means per generation. Here, one can see that GA is indeed performing better than DAPTEV in terms of docking score statistics across generations.

**Figure 6.**
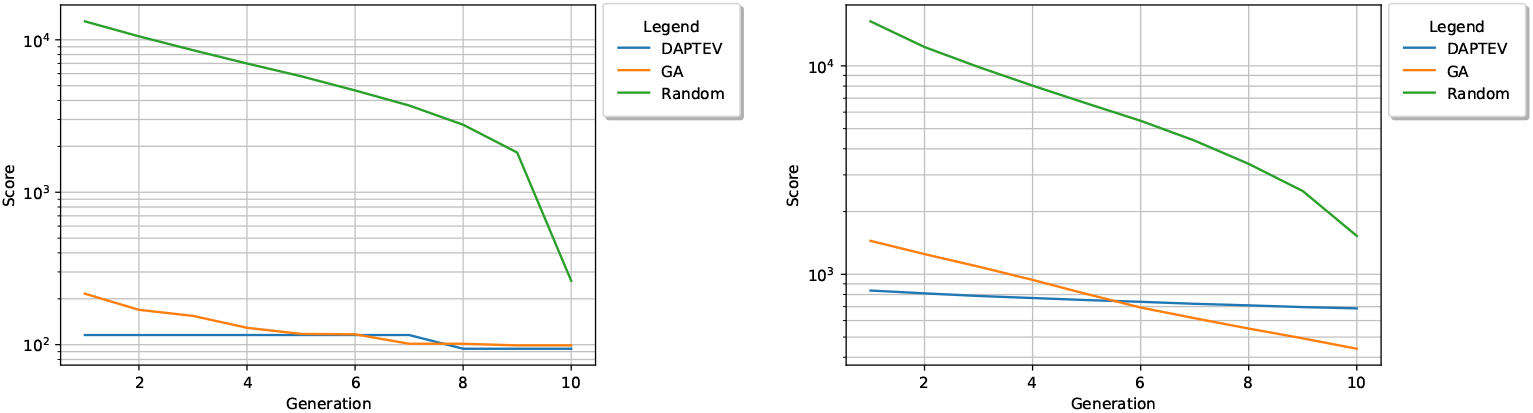
The three models’ 5 best (left) and worst (right) mean scores plotted per generation on a logarithmic scale.

Furthermore, to have a full view of the binding score distribution for each method, the density curves of Aptamers’ binding scores in the last generations of DAPTEV, GA, and HC (i.e., Random) are displayed in Fig. 7. From the density plots, one can conclude that (1) DAPTEV and GA obtained Aptamer sequences with significantly better binding scores than HC (Mann-Whitney U test, p-value=7.25e-226 and p-value=2.31e-256, respectively), and (2) GA’s result is better than DAPTEV’s result in terms of docking score distribution (Mann-Whitney U test, p-value=8.47e-142).

**Figure 7.**
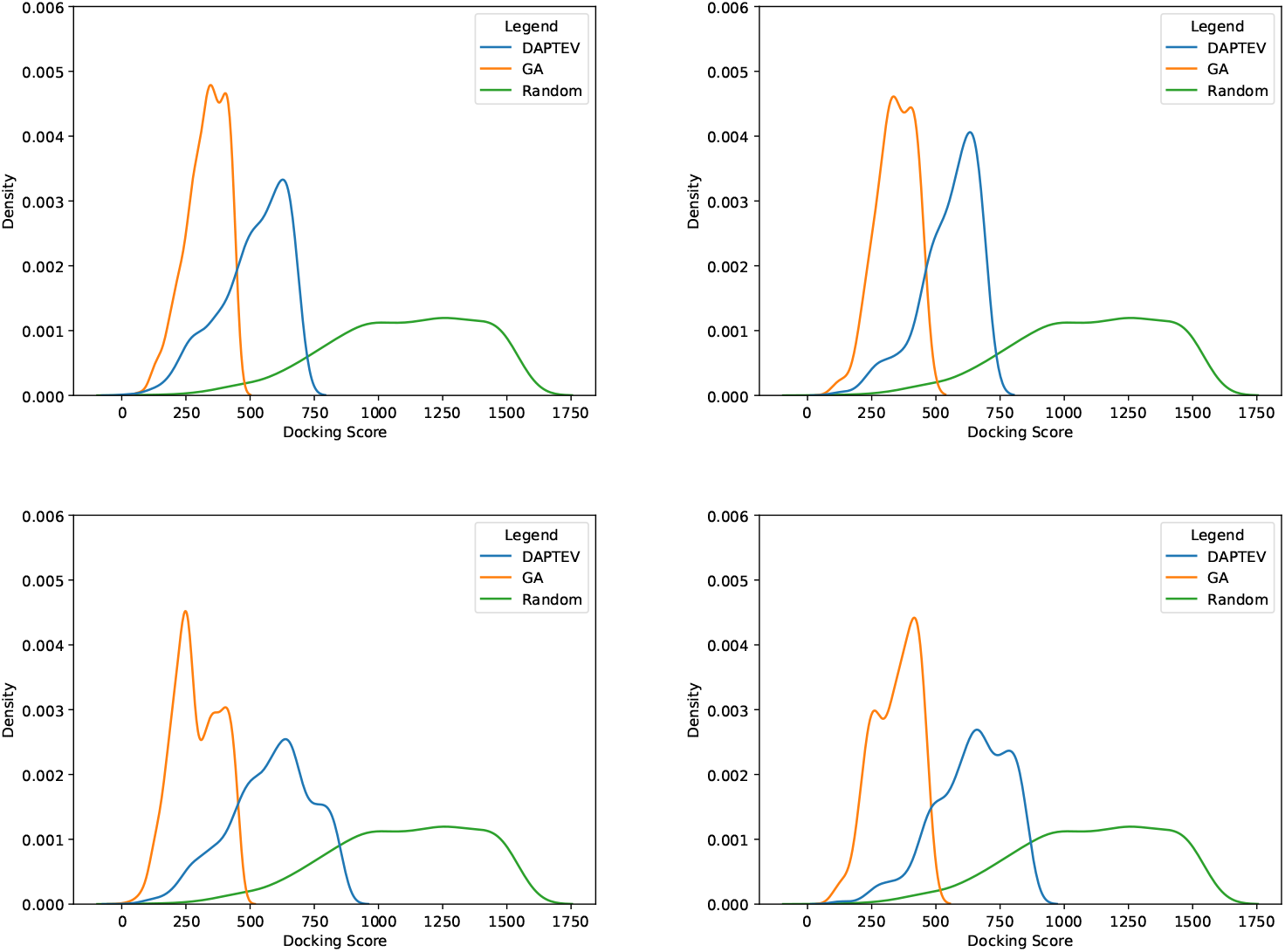
Density curves of binding scores of aptamer samples in the last generations of DAPTEV, GA, and HC (Random), respectively. The top-left graph shows all scores produced in run 2. The top-right graph shows all folded complex scores from run 2. The bottom-left shows all scores from all three runs. The bottom-right shows all folded complex scores from all three runs.

### Comparison in Terms of Base Pairing

Thus far, it has seemed as though GA is the best suited model for this application. However, there are two goals for this experiment. While the first goal is to optimize the docking scores, the second goal is to produce sequences with at least one connection (nucleotide base pairing) in the secondary structures. In other words, these models were also supposed to learn the key features of well-performing structural motifs applied to the SARS-CoV-2 spike protein RBD.

Table 3 provides further details on the produced folded sequences. The GA performed rather poorly in this task, only producing 47 (on average) folded structures out of 800 total sequences, achieving an 6% fold rate. Conversely, DAPTEV produced 272 (on average) folded structures out of 800 total sequences, achieving a 34% fold rate. It is important to note that the HC (random) algorithm entirely constructed folded RNAs as the dataset creation script does not allow for unfolded structures in its random search.

**Table 3.**
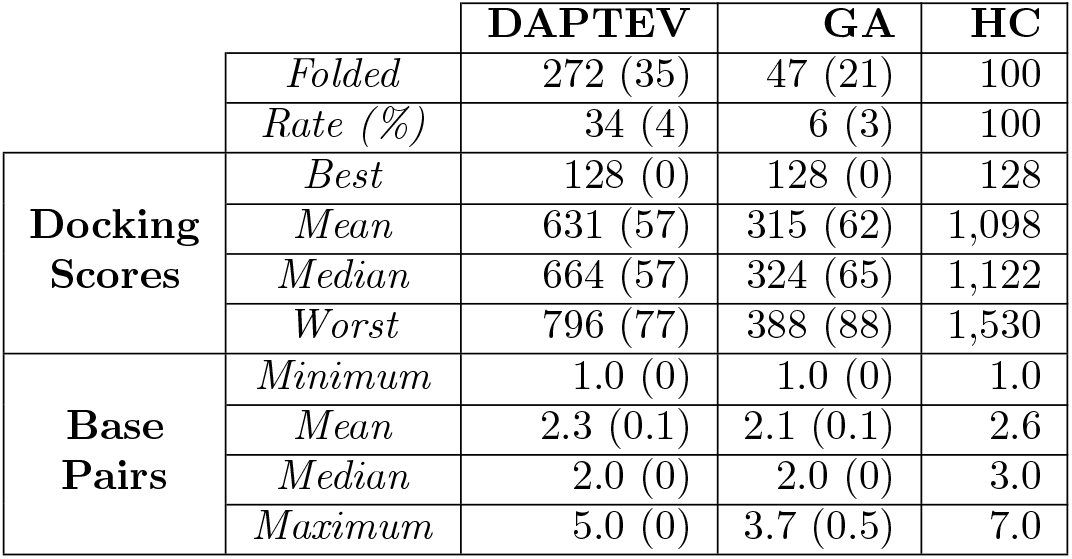
Average docking scores and base pairings relating to the folded RNAs in each model. Standard deviation provided in brackets. Note: all values relate only to the folded RNAs in the last generation.

In Fig. 8a, the best folded complex has a docking score of 128 with two base pairings in its secondary structure. Interestingly, every model produced the same best performing folded RNA. The median complex has a score of 653, also with two base pairings, and the worst has a score of 857 with one base pairing. Fig. 8b has a median docking score of 358 with four base pairings, and the worst score of 460 has three base pairings. Every complex in Fig. 8c has folded RNAs due to the nature of the dataset creation script. The median score is 1,122 with three base pairings and the worst score is 1,530 with one base pairing.

**Figure 8.**
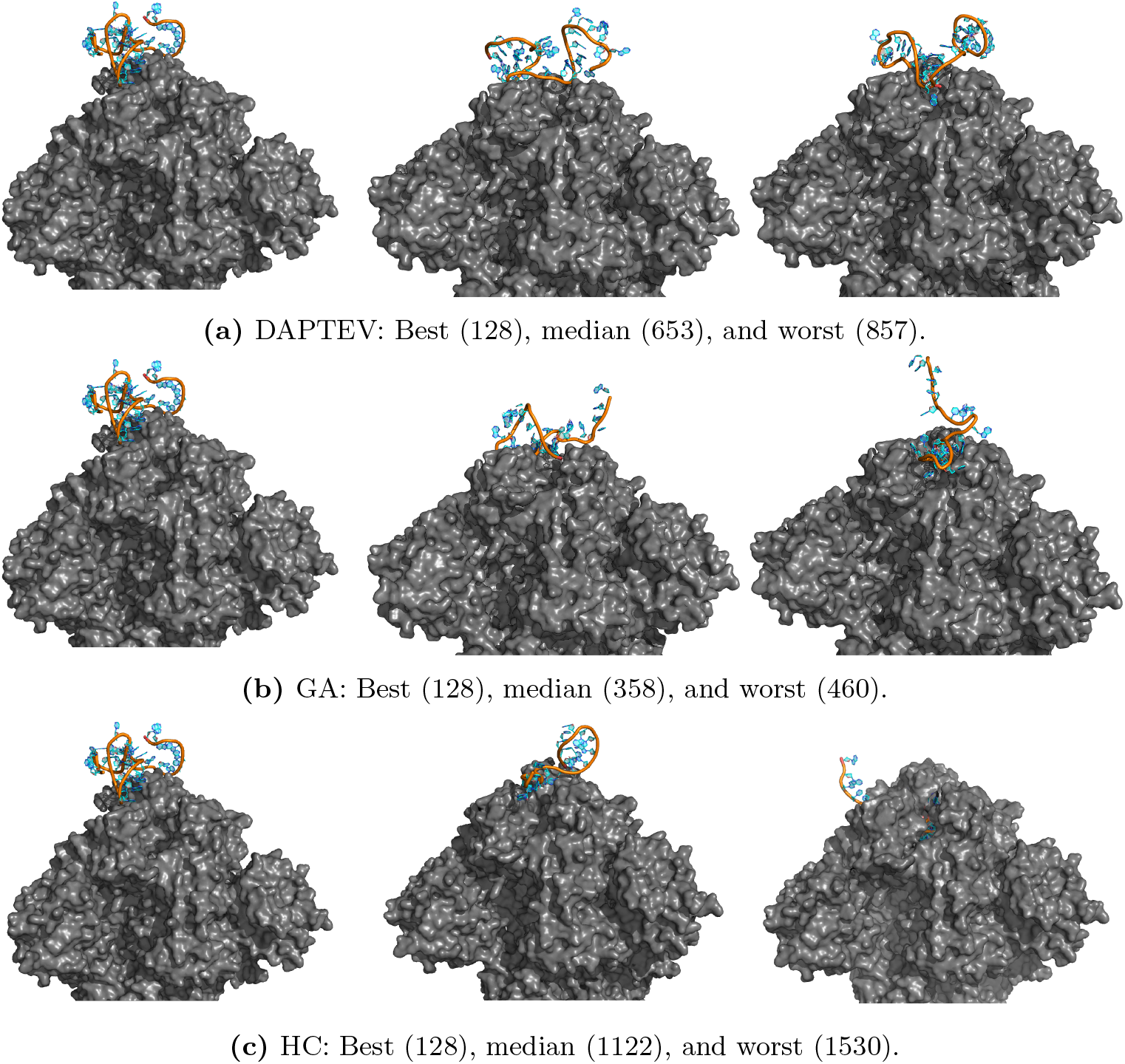
Folded RNAs docked to the SARS-CoV-2 RBD based on Rosetta-returned scores. Associated docking scores are in brackets next to the classification of each complex.

### Novelty and Diversity of Generated Aptamers

Lastly, two additional metrics were calculated to assess model performance. These metrics, expressed as ratios, include novelty and diversity. *Novelty* is calculated as the number of generated sequences that do not exist in the initial training data versus the total number of generated sequences in a generation (the “population size”). *Diversity* is calculated as the number of generated unique sequences versus the total number of generated sequences in a generation. The novelty values calculated for DAPTEV and GA’s last generations are 0.8475 and 0.9962 respectively. Similarly, the diversity values were 0.8487 and 0.9937 respectively. Hence, both DAPTEV and GA show good novelty and diversity, and GA seems always to produce novel and unique sequences.

## Discussion and Conclusion

### Observations

While it may seem that GA performed the best in terms of docking scores, this is not actually the case if the aptamer structures are considered as well. As previously mentioned, the docking score can be artificially improved by producing an unfolded RNA secondary structure which, in turn, will incur fewer penalties during the RNA tertiary structure prediction and the docking simulation. If a model prioritizes only unconnected structures, it stands to reason that it would seem to perform better when only considering score output. However, the expectation of this research is three-fold. (1) The docking scores has to be optimized. This was accomplished best by GA as substantiated by the returned scores and statistical analysis. (2) Some learning of well-performing structural motifs in the provided RNA secondary structures is required.Producing unfolded RNAs is not overly helpful when attempting to develop aptamer-based drugs. Even if this is a desirable trait, we expect to produce more complex structures from intelligent models and not be forced to sacrifice these features. Based on the percentage of folded structures in the last generation, it is clear that DAPTEV performs significantly better than GA. This points to the conclusion that GA solely prioritized the optimization of scores to the detriment of structural complexity. These results also indicate that DAPTEV is indeed able to learn structural motif patterns from the training data. (3) The deep learning model has to possess the ability to be queried for new, well-performing, structurally connected RNAs to explore aptamer-based drug discovery. This point implies the persistence of a trained model and a way to request new data. A GA must be trained every time new sequences are required. A VAE, conversely, can have its current state of learning saved and reloaded nearly immediately. Furthermore, new sequences can be obtained immediately. With everything mentioned above, one could posit that DAPTEV performs admirably in all three requirements. Additionally, every score produced in the last generation of DAPTEV was below the score threshold of 3,500, which was docking score limit set initially to tell the VAE that a sequence was performing well. DAPTEV was able to learn structural features and other characteristics that make RNA perform well when docking to a specific target. Finally, the model can be saved and queried for new sequences that should perform well at docking to a specific target’s RBD while also maintaining some structural complexity. In contrast, GA and the hill climber algorithm must both be retrained every time, GA prioritized score optimization at the cost of motif preservation, and the hill climber is likely to yield poor results due to premature convergence at a local minimum.

### Limitations

There are some limitations of this research. Firstly, DAPTEV did not have enough starting aptamer data in the initial dataset. Only 849 unique, known, aptamer sequences were found during the literature review stage, but that was narrowed down further to only 344 due to parameter and computation time constraints.

Secondly, at present DAPTEV does not work with DNAs due to the limitations of utilized software. While this research was mainly focused on RNAs, some other research has worked on DNAs. It would have been preferable to include DNAs in the capabilities of this model.

The randomly generated sequences must have at least one connection. However, more control over the secondary structure is preferred. The specification of connection amounts or a range of acceptable connections would be beneficial. Furthermore, the DAPTEV model is affected by limited accuracy from the structure/energy prediction process of both secondary and tertiary structures.

Currently, DAPTEV assumes that the latent space follows a Gaussian distribution. This distribution may not be robust enough to capture the complexity of the task. Testing different distributions could have yielded further insight into the VAE’s ability to operate in this problem space.

### Conclusion

The goal of this research was to see if a deep generative model would be efficient at accelerating the RNA aptamer drug development process. While this research was applied to the SARS-CoV-2 spike protein, careful consideration was placed into the universal design for nearly any protein target. With regard to target affinity, one could conclude that both DAPTEV and GA performed well at this task. While the GA did outperform DAPTEV in this regard, the difference between these two models was not very large. Especially when considering that the score threshold was set to 3,500 and DAPTEV was still able to produce scores significantly lower than that. For target specificity, DAPTEV certainly shows some promising results. This is especially true when compared to GA. DAPTEV had an output with 34% of its produced sequences containing at least one connection in the secondary structure. This number could have been even higher if more vocabulary combinations had been implemented and some additional parameter tests were run. Conversely, GA only produced a 6% fold rate. This indicates that the VAE was indeed able to learn some structural features of the data and the MLP regularizer performed well at its task. The code for this model is accessible at https://github.com/candress/DAPTEV_Model.

### Future Work

Several additional features and considerations could improve this research. (1) One such consideration is the fact that there was not enough starting real aptamer data. It would have been preferable to add more to the starting dataset. Unfortunately, at the time of searching, it seemed most of the aptamer datasets and databases were taken down. (2) Currently, this model only accepts RNA data. However, DNA aptamers are more common in literature. It would be beneficial to include the capability to work with DNA data. (3) Allowing the user to perform tertiary structure prediction separately, or to provide pre-calculated tertiary structure could be beneficial. (4) Also, statically choosing the protein atom of CA and the RNA atom of C5 may be negatively affecting the performance as there is no determination of which atoms are more likely to interact. Instead, the interactions between each atom present in the protein residue and the possible RNA bases could have been pre-calculated. (5) It would be beneficial to increase the vocabulary combination amount. This DAPTEV only allows for combinations up to three characters long to reduce computation time. However, a vocabulary length of size three impedes the model’s ability to learn more complex structural motifs. (6) Some additional considerations and implementations could be the following: a learnable KL/reconstruction loss balancing component could be implemented so the model regulates the weights of the KL and the reconstruction loss [53]. Transformer is another versatile deep learning model that relies entirely on self-attention and is well-suited for design tasks [54], but requires more data to train. Given more data available in the future, it would be interesting to switch out the VAE with a Transformer to see how the results are affected. (7) It is difficult to make conclusive observations at this moment due to the few number of experimental runs. Ideally, at least 30 runs per experiment per model should be performed. However, this would take an inordinate amount of time given the current computing restrictions for this research. The only way this would be feasible is if one had access to a cluster to distribute the workload. (8) Our discovered new aptamers with promising binding affinity and specificity to the SARS-CoV-2 spike protein can be tested in wet-lab experiments.

## Acknowledgements

The use of Rosetta was technical supported by Das Lab’s Dr. Rhiju Das, Dr. Ramya Rangan, and Dr. Andy Watkins. Funding was provided by the AI for Design Challenging Program at the National Research Council Canada, the Discovery Grant from the Natural Sciences and Engineering Research Council of Canada, and the Ontario Graduate Scholarships. Dr. Kalli Kappel is supported by the Schmidt Science Fellows, in partnership with the Rhodes Trust, and the HHMI Hanna H. Gray Fellows Program. Additional insights were rendered by Brock University’s Dr. Robson De Grande, Dr. Sheridan Houghten, and Dr. Ali Emami.

## References

1. Kim TH, Lee SW. Aptamers for Anti-Viral Therapeutics and Diagnostics. International Journal of Molecular Sciences. 2021;22(8):4168.

2. Lan J, Ge J, Yu J, Shan S, Zhou H, Fan S, et al. Structure of the SARS-COV-2 Spike Receptor-binding Domain Bound to the ACE2 Receptor. Nature. 2020;581(7807):215–220.

3. Song Y, Song J, Wei X, Huang M, Sun M, Zhu L, et al. Discovery of Aptamers Targeting the Receptor-Binding Domain of the SARS-CoV-2 Spike Glycoprotein. Analytical Chemistry. 2020;92(14):9895–9900.

4. Tai W, He L, Zhang X, Pu J, Voronin D, Jiang S, et al. Characterization of the Receptor-binding Domain (RBD) of 2019 Novel Coronavirus: Implication for Development of RBD Protein as a Viral Attachment Inhibitor and Vaccine. Cellular and Molecular Immunology. 2020;17(6):613–620.

5. Walls AC, Park YJ, Tortorici MA, Wall A, McGuire AT, Veesler D. Structure, Function, and Antigenicity of the SARS-CoV-2 Spike Glycoprotein. Cell. 2020;181(2):281–292.e6.

6. Yi C, Sun X, Ye J, Ding L, Liu M, Yang Z, et al. Key Residues of the Receptor Binding Motif in the Spike Protein of SARS-CoV-2 that Interact with ACE2 and Neutralizing Antibodies. Cellular & Molecular Immunology. 2020;17(6):621–630.

7. Wengerter BC, Katakowski JA, Rosenberg JM, Park CG, Almo SC, Palliser D, et al. Aptamer-targeted Antigen Delivery. Molecular Therapy: The Journal of the American Society of Gene Therapy. 2014;22(7):1375–1387.

8. Yoon SY, Gee G, Hong KJ, Seo SH. Application of Aptamers for Assessment of Vaccine Efficacy. Clinical and Experimental Vaccine Research. 2017;6(2):160–163.

9. Plotkin S, Robinson JM, Cunningham G, Iqbal R, Larsen S. The Complexity and Cost of Vaccine Manufacturing – An Overview. Vaccine. 2017;35(33):4064–4071.

10. Tuerk C, Gold L. Systematic Evolution of Ligands by Exponential Enrichment: RNA Ligands to Bacteriophage T4 DNA Polymerase. Science. 1990;249(4968):505–510.

11. Kupakuwana GV, II JEC, McPike MP, Borer PN. Acyclic Identification of Aptamers for Human Alpha-thrombin Using Over-represented Libraries and Deep Sequencing. PloS One. 2020;6(5):e19395.

12. Ahirwar R, Nahar S, Aggarwal S, Ramachandran S, Maiti S, Nahar P. In Silico Selection Of An Aptamer To Estrogen Receptor Alpha Using Computational Docking Employing Estrogen Response Elements As Aptamer-Alike Molecules. Scientific Reports. 2016;6(1):21285.

13. Lee G, Jang GH, Kang HY, Song G. Predicting Aptamer Sequences that Interact with Target Proteins Using an Aptamer-protein Interaction Classifier and a Monte Carlo Tree Search Approach. PLOS ONE. 2021;16(6):e0253760.

14. Song J, Zheng Y, Huang M, Wu L, Wang W, Zhu Z, et al. A Sequential Multidimensional Analysis Algorithm for Aptamer Identification Based on Structure Analysis and Machine Learning. Analytical Chemistry. 2020;92(4):3307–3314.

15. Becker KCD, Becker RC. Nucleic Acid Aptamers as Adjuncts to Vaccine Development. Current Opinion in Molecular Therapeutics. 2006;8(2):122–129.

16. Chen Z, Hu L, Zhang BT, Lu A, Wang Y, Yu Y, et al. Artificial Intelligence in Aptamer–Target Binding Prediction. International Journal of Molecular Sciences. 2021;22(7):3605.

17. Keefe AD, Pai S, Ellington A. Aptamers as Therapeutics. Nature Reviews Drug Discovery. 2010;9(7):537–550.

18. Kinghorn AB, Fraser LA, Lang S, Shiu SCC, Tanner JA. Aptamer Bioinformatics. International Journal of Molecular Sciences. 2017;18(12).

19. Wornow M. Applying Deep Learning to Discover Highly Functionalized Nucleic Acid Polymers That Bind to Small Molecules [Bachelor’s Thesis]. Harvard College. Cambridge, USA; 2020.

20. Zou X, Wu J, Gu J, Shen L, Mao L. Application of Aptamers in Virus Detection and Antiviral Therapy. Frontiers in Microbiology. 2019;10:1462.

21. Russell S, Norvig P. Artificial Intelligence: A Modern Approach. 4th ed. Pearson; 2020.

22. Bishop CM. Pattern Recognition and Machine Learning. New York: Springer; 2006.

23. LeCun Y, Bengio Y, Hinton G. Deep Learning. Nature. 2015;521:436–444.

24. Romez-Bombarelli R, Wei JN, Duvenaud D, Hernarndez-Lobato JM, Sanchez-Lengeling B, Sheberla D, et al. Automatic Chemical Design Using a Data-driven Continuous Representation of Molecules. ACS Central Science. 2018;4:268–276.

25. Kim J, Park S, Min D, Kim W. Comprehensive Survey of Recent Drug Discovery Using Deep Learning. International Journal of Molecular Sciences. 2021;22:9983.

26. Grantham K, Mukaidaisi M, Ooi HK, Ghaemi MS, Tchagang A, Li Y. Deep Evolutionary Learning for Molecular Design. IEEE Computational Intelligence Magazine. 2022;17(2):14–28.

27. Mukaidaisi M, Vu A, Grantham K, Tchagang A, Li Y. Multi-objective Drug Design Based on Graph-fragment Molecular Representation and Deep Evolutionary Learning. Frontier in Pharmacology. 2022;13:920747.

28. Jiang P, Meyer S, Hou Z, Propson NE, Soh HT, Thomson JA, et al. MPBind: A Meta-motif-based Statistical Framework and Pipeline to Predict Binding Potential of SELEX-derived Aptamers. Bioinformatics. 2014;30(18):2665–2667.

29. Hoinka J, Berezhnoy A, Sauna ZE, Gilboa E, Przytycka TM. AptaCluster - A Method to Cluster HT-SELEX Aptamer Pools and Lessons from Its Application. In: International Conference on Research in Computational Molecular Biology; 2014. p. 115–128.

30. Alam KK, Chang JL, Burke DH. FASTAptamer: A Bioinformatic Toolkit for High-throughput Sequence Analysis of Combinatorial Selections. Molecular Therapy - Nucleic Acids. 2015;4:e230.

31. Hiller M, Pudimat R, Busch A, Backofen R. Using RNA Secondary Structures to Guide Sequence Motif Finding Towards Single-stranded Regions. Nucleic Acids Research. 2006;34(17):e117.

32. Hoinka J, Zotenko E, Friedman A, Sauna ZE, Przytycka TM. Identification of Sequence-structure RNA Binding Motifs for SELEX-derived Aptamers. Bioinformatics. 2012;28(12):i215–i223.

33. Dao P, Hoinka J, Takahashi M, Zhou J, Ho M, Wang Y, et al. AptaTRACE Elucidates RNA Sequence-Structure Motifs from Selection Trends in HT-SELEX Experiments. Cell Systems. 2016;3:62–70.

34. Li BQ, Zhang YC, Huang GH, Cui WR, Zhang N, Cai YD. Prediction of Aptamer-Target Interacting Pairs with Pseudo-Amino Acid Composition. PLoS ONE. 2014;9(1):e86729.

35. Zhang L, Zhang C, Gao R, Yang R, Song Q. Prediction of Aptamer-protein Interacting Pairs Using an Ensemble Classifier in Combination with Various Protein Sequence Attributes. BMC Bioinformatics. 2016;17:225.

36. Peng C, Han S, Zhang H, Li Y. RPITER: A Hierarchical Deep Learning Framework for ncRNA–Protein Interaction Prediction. International Journal of Molecular Sciences. 2019;20:1070.

37. Im J, Park B, Han K. A Generative Model for Constructing Nucleic Acid Sequences Binding to a Protein. BMC Genomics. 2019;20(13):967.

38. Alipanhi B, Delong A, Weirauch MT, Frey BJ. Predicting the Sequence Specificities of DNA-and RNA-binding Proteins by Deep Learning. Nature Biotechnology. 2015;33(8):831–838.

39. Park B, Han K. Discovering Protein-binding RNA Motifs with a Generative Model of RNA Sequences. Computational Biology and Chemistry. 2020;84.

40. Iwano N, Adachi T, Aoki K, Nakamura Y, Hamada M. RaptGen: A Variational Autoencoder with Profile Hidden Markov Model for Generative Aptamer Discovery. bioRxiv. 2021; p. 2021.02.17.431338.

41. Chaslot GMJB, Winands MHM, Herik HJVD, Uiterwijk JWHM. Progressive Strategies for Monte-Carlo Tree Search. New Mathematics and Natural Computation. 2008;4(3):343–357.

42. Kappel K, Das R. Sampling Native-like Structures of RNA-Protein Complexes Through Rosetta Folding and Docking. Structure. 2019;27(1):140–151.e5.

43. Wayment-Steele HK, Kladwang W, Strom AI, Lee J, Treuille A, Eterna Participants, et al. RNA Secondary Structure Packages Evaluated and Improved by High-throughput Experiments. Biophysics; 2020.

44. Kingma DP, Welling M. Auto-Encoding Variational Bayes. In: International Conference on Learning Representations; 2014.

45. Chung J, Gulcehre C, Cho K, Bengio Y. Empirical Evaluation of Gated Recurrent Neural Networks on Sequence Modeling. In: NIPS 2014 Deep Learning and Representation Learning Workshop; 2014.

46. Mikolov T, Chen K, Corrado GS, Dean J. Efficient Estimation of Word Representations in Vector Space. In: International Conference on Learning Representations; 2013.

47. Pennington J, Socher R, Manning C. GloVe: Global Vectors for Word Representation. In: Conference on Empirical Methods in Natural Language. Doha, Qatar: Association for Computational Linguistics; 2014. p. 1532–1543.

48. Fu H, Li C, Liu X, Gao J, Celikyilmaz A, Carin L. Cyclical Annealing Schedule: A Simple Approach to Mitigating KL Vanishing. ArXiv. 2019; p. arXiv.1903.10145.

49. Premkumar L, Segovia-Chumbez B, Jadi R, Martinez DR, Raut R, Markmann A, et al. The Receptor Binding Domain of the Viral Spike Protein is an Immunodominant and Highly Specific Target of Antibodies in SARS-CoV-2 Patients. Science Immunology. 2020;5(48):eabc8413.

50. Markley JL, Bax A, Arata Y, Hilbers CW, Kaptein R, Sykes BD, et al. Recommendations for the Presentation of NMR Structures of Proteins and Nucleic Acids – IUPAC-IUBMB-IUPAB Inter-Union Task Group on the Standardization of Data Bases of Protein and Nucleic Acid Structures Determined by NMR Spectroscopy. Journal of Biomolecular NMR. 1998;12(1):1–23.

51. Berman HM, Westbrook J, Feng Z, Gilliland G, Bhat TN, Weissig H, et al. The Protein Data Bank. Nucleic Acids Research. 2000;28(1):235–242.

52. Li BQ, Zhang YC, Huang GH, Cui WR, Zhang N, Cai YD. Prediction of Aptamer-Target Interacting Pairs with Pseudo-Amino Acid Composition. PLOS ONE. 2014;9(1):e86729.

53. Asperti A, Trentin M. Balancing Reconstruction Error and Kullback-Leibler Divergence in Variational Autoencoders. arXiv. 2020; p. ArXiv.2002.07514.

54. Vaswani A, Shazeer N, Parmar N, Uszkoreit J, Jones L, Gomez AN, et al. Attention is All You Need. In: Neural Information Processing Systems; 2017.

